# Astronauts Gaze Behavior in Visual Target Acquisition during Space Shuttle Flights

**DOI:** 10.1101/2022.05.25.493475

**Authors:** Ognyan I. Kolev, Millard F. Reschke, Gilles Clément

## Abstract

We explored how weightlessness during space flight altered the astronauts’ gaze behavior with respect to flight day and target eccentricity. Thirty-four astronauts of 20 Space Shuttle missions had to acquire visual targets with angular offsets of 20°, 30°, and 49°. Measurements of eye, head, and gaze positions collected before and during flight days 1 to 15 indicated changes during target acquisition that varied as a function of flight days and target eccentricity. The in-flight changes in gaze behavior were presumably the result of a combination of several factors, including a transfer from allocentric to egocentric reference for spatial orientation in absence of a gravitational reference, the generation of slower head movements to prevent space motion sickness, and a decrease in smooth pursuit and vestibulo-ocular reflex performance. These results confirm that humans have several strategies for gaze behavior, between which they switch depending on the environmental conditions.

## INTRODUCTION

Coordination of motor activity is adapted to Earth’s gravity (1 g). However, during space flight the gravity level changes from Earth gravity to hypergravity during launch, and to microgravity (0 g) in orbit. This transition between gravity levels may alter the coordination between eye and head movements in gaze performance for visual target acquisition, which is critical for piloting and controlling onboard instruments.

Previous experiments on visual target acquisition during space flight have given conflicting results. Saccadic peak eye velocity has been reported to increase in some astronauts, and to decrease in others. Also, the amplitude of eye movements during horizontal and vertical tracking increased in orbit in some cosmonauts, but decreased in others (Vesterhauge et al. 1984, André-Deshays et al. 1993, Kornilova et al. 1993, Uri et al. 1989, Somers et al. 2002). One problem when comparing the results across these previous studies is that measurements were obtained on different flight days (FD). Also, the first in-flight measurements of eye-head coordination in these studies were made from flight day 5 (André-Deshays et al. 1993), so the immediate effects of exposure to 0 g were not documented.

The purpose of the present study was to explore whether and how entry into weightlessness altered gaze behavior for visual target acquisition. We report here the results obtained in 34 astronauts starting on FD1 and up to FD15. Because of the changes in vestibular and proprioceptive signals sensory, perceived space orientation, background brain activity, and biomechanics of head motion immediately after insertion into orbit, we hypothesized that eye-head coordination during visual target acquisition would be altered in 0 g compared to 1 g, and this alteration would decrease over a period of two week in space.

## MATERIALS AND METHODS

### Subjects

Thirty-four NASA astronauts (30 males and 4 females) ranging in age from 34 to 54 years (mean 41 years) participated in this investigation. These astronauts flew on board 20 Space Shuttle missions. None of the participants had any ocular motor, vestibular abnormalities, or were taking any drugs with effects on the nervous system. The experimental protocol was approved by the NASA Johnson Space Center Institutional Review Board. All astronauts gave their written informed consent to participate in this investigation in accordance with the tenets of the Declaration of Helsinki.

### Visual Targets

Visual targets subtending 0.5° were fixed to a screen at a distance of 86.4 cm from the subjects. Targets were located at angular distances of 20°, 30°, and 49° off center in the horizontal plane at subject’s eye level. To easily differentiate between targets, each target corresponding to its degree of angular offset from center was color-coded (20° green, 30° red, 49° blue).

### Eye and Head Measurements

Horizontal eye positions were measured using standard electro-oculography (EOG). Two electrodes were applied to the outer canthus of each eye, and one ground electrode was applied to a neutral surface behind the right ear. The EOG signals were amplified and digitized at a sampling rate of 500 Hz. Extraneous high frequency noise was digitally removed by filtering before processing with a finite impulse response low-pass Hamming window filter with a cutoff frequency of 30 Hz.

Horizontal head velocities were measured using a tri-axial angular rate sensor fixed to the head with the webbing from a hardhat liner. The angular rate sensor was located approximately on the apex of the skull and adjusted prior to each test session to minimize cross talk between all 3 axes. The signals from the rate sensor were digitized at 500 Hz and filtered using the same procedure as the EOG signals.

Horizontal head position was calibrated using a low power laser mounted on the hardhat liner that could be adjusted so that the laser was located centrally on the forehead between the two eyes. When activated, the subject moved the laser point on the central target and to each of the color-coded targets.

### Experimental Protocol

Eye movement calibration was performed by having the subjects oscillating their heads at approximately 0.25 Hz at amplitude of 30° to the right and to the left while fixating the central target. Based on the vestibulo-ocular reflex (VOR) thus generated, the calculated angular head position was used to determine the expected eye position required to maintain visual fixation. These expected eye positions were compared with the corresponding EOG signals to yield the volts-to-degree relationship necessary for calculating a calibration curve. Eye velocity was obtained through differentiation of the filtered eye position signal.

During the visual target acquisition task, the operator called out the various targets (“right green”, “left green”, “right red”, “left red”, “right blue”, “left blue”) in a predetermined random order. The crewmembers were asked to look from a central fixation point to the called target as quickly and accurately as possible using both the head and eyes. Crewmembers maintained fixation on the called target for a minimum of 1.5 s, and then returned their gaze to the center target for approximately 10 s. This was to ensure that the cupula of the horizontal semicircular canals had returned to its original position before the next trial. Each target was tested between 2-5 trials depending on time constraints.

For the 20° and 30° targets, data was collected 10 days before launch (Pre, n=34) and then on FD1 (n=32), FD2 (n=33), FD5 (n=20), and FD15 (n=6). A subset of our astronaut subjects performed the experiment for the 49° target. This condition was considered a supplemental objective, and unfortunately no preflight data was collected for this target eccentricity. Nevertheless, in orbit, the experiment with the 49° target was performed on FD1 (n=12), FD2 (n=10), FD3 (n=3), FD5 (n=11), FD7 (n=9), FD11 (n=13), FD13 (n=2), and FD15 (n=6).

Figure 1 represents a typical target acquisition trial. Gaze is the sum of head and eye positions. The following duration, position, and velocity parameters were calculated from the eye, head, and gaze measurements: (a) time to primary saccade; (b) duration of primary saccade; (c) amplitude of primary saccade; (d) peak velocity of primary saccade; (e) time to final position; and (f) final position. The peak acceleration of the primary head saccade was also calculated.

**Figure 1.**
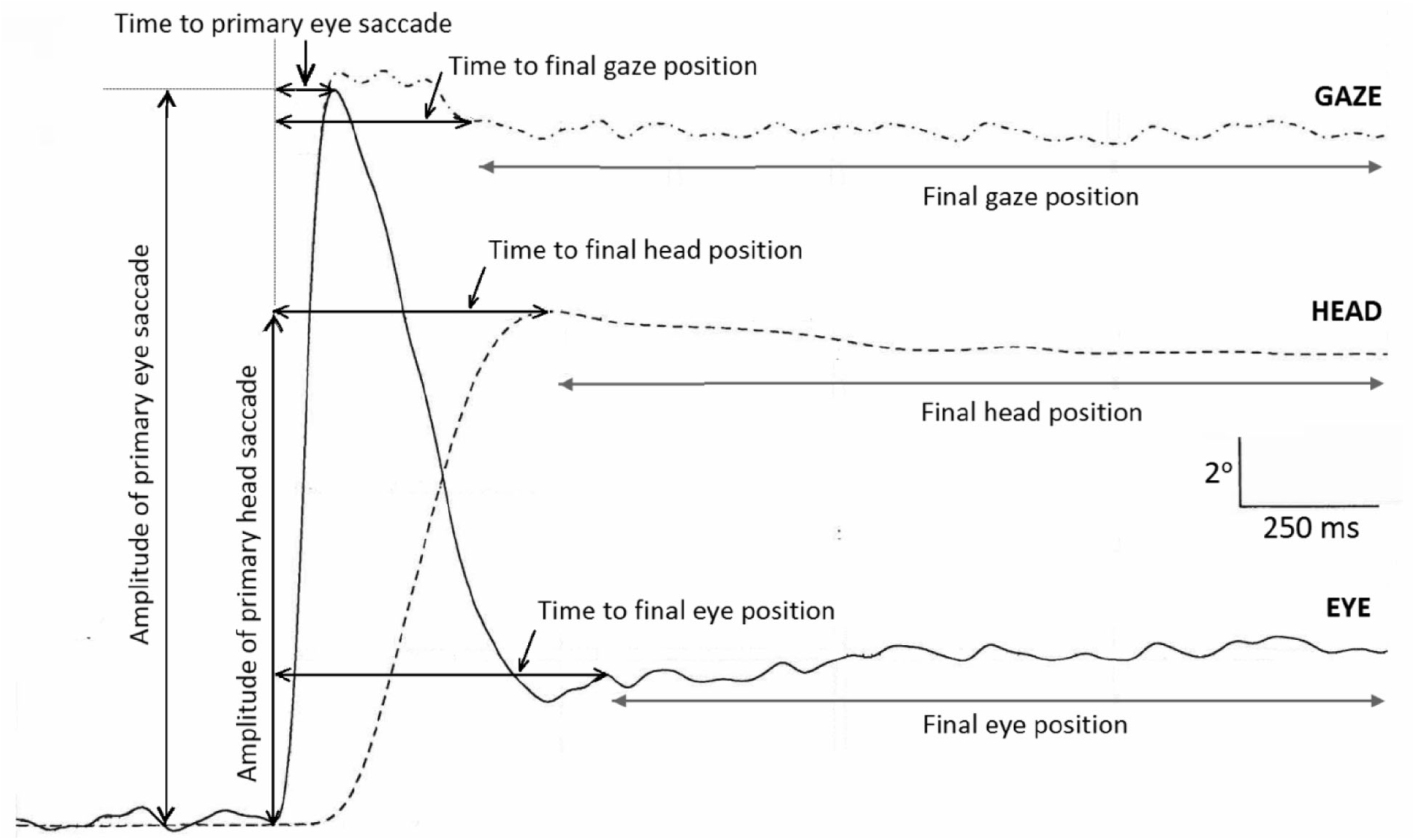
Time sequence of eye and head, and gaze (sum of eye and head) during visual acquisition of a 30° target.

### Statistical Analysis

Two-way (session day: Pre, FD1, FD2, FD5, FD15; target offset: 20°, 30°) ANOVAs were used. The measurement prior flight was taken only at 20° and 30° offsets. One-way ANOVAs were used for the 49° target offset with factor days: FD1, FD2, FD3, FD5, FD7, FD11, FD13, and FD15.

After the completion of the ANOVAs, Fisher Least Significant Difference (LSD) tests were used for pairwise comparisons of the means, valid only for the significant effects. A plot of the residuals versus fitted values and the quantiles plots (Normal and Half-Normal) was used for each response variable to demonstrate the adequacy of the distributional assumptions regarding use of linear regressions and ANOVAs.

All analyses were conducted using statistical package GenStat (VSN Int, 2017), R environment (R Core Team, 2016), and R package ggplot2.

## RESULTS

### Eye and Head Coordination Strategy

During the acquisition of visual targets situated 20° off-center and more, humans typically use both eye and head movements. The closer the target is to the extreme oculomotor range (about 55°), the more the head contributes to the performance (Guitton and Volle 1987). The head, having greater inertia than the eye, typically moves after the eye has moved in the orbit. At the end of the eye saccade the gaze has already reached its final position, but the head continues to move. The head movement stimulates the horizontal semicircular canals, which produces an eye movement through the VOR that is opposite in direction and velocity to that of the head. When moving to a target gaze often undershoots or overshoots the target and corrective eye saccades occur to reposition the gaze on the target. In general, the total gaze movement is greater than the total head movement, so that the final position of the eye is offset in the direction of head movement (Reschke et al. 2017). The results from the statistical analysis using ANOVAs are presented in Tables 1 and 2.

**Table 1.** Statistical significance (P-value) of the main effects and the interaction of factors ‘day’ and ‘target’ for head, gaze, and eye during two-way ANOVA’s for the 20°, 30°, and 49°targets. NS: not significant; NA: not applicable.

| Response Variable | Factors | Gaze | Head | Eye |
| --- | --- | --- | --- | --- |
| *Time to Final Position* | day | <.001 | <.001 | <.001 |
|  | target | <.001 | 0.012 | 0.002 |
|  | day x target | 0.074 | <.001 | 0.097 |
| *Final Position* | day | <.001 | <.001 | <.001 |
|  | target | <.001 | <.001 | <.001 |
|  | day x target | <.001 | 0.048 | <.001 |
| *Time to Primary Saccade* | day | <.001 | <.001 | <.001 |
|  | target | <.001 | <.001 | <.001 |
|  | day x target | 0.002 | 0.009 | <.001 |
| *Amplitude of Primary Saccade* | day | <.001 | <.001 | <.001 |
|  | target | <.001 | <.001 | <.001 |
|  | day x target | <.001 | 0.161 | 0.027 |
| *Peak Velocity of Primary Saccade* | day | <.001 | NS | <.001 |
|  | target | <.001 | <.001 | NS |
|  | day x target | NS | NS | NS |
| *Duration of Primary Eye Saccade* | day | NA | NA | <.001 |
|  | target | NA | NA | <.001 |
|  | day x target | NA | NA | 0.034 |
| *Acceleration of Primary Head Saccade* | day | NA | 0.018 | NA |
|  | target | NA | <.001 | NA |
|  | day x target | NA | 0.018 | NA |

**Table 2.** Statistical significance (P-value) of the main effects of factor day for gaze, head, and eye. Results of one-way ANOVA for 49° targets. NS: not significant; NA: not applicable.

| **Response Variable** | **Factor** | **Gaze** | **Head** | **Eye** |
| --- | --- | --- | --- | --- |
| *Time to Final Position* | day | .007 | <.001 | <.001 |
| *Final Position* | day | NS | <.001 | <.001 |
| *Time to Primary Saccade* | day | <.001 | <.001 | <.001 |
| *Amplitude of Primary Saccade* | day | <.001 | <.001 | <.001 |
| *Peak Velocity of Primary Saccade* | day | <.001 | 0.036 | <.001 |
| *Duration of Primary Eye Saccade* | day | NA | NA | <.001 |
| *Acceleration of Primary Head Saccade* | day | NA | <.001 | NA |

### Gaze Dynamics

The final position of gaze was right on target for all preflight and in-flight trials (Figure 2A). However, the time to reach this final position was different across flight days (Figure 2B). This duration was longer than preflight with a peak on FD2 for the 20° and 30° targets. On FD15 the time for the gaze to reach the target had returned to baseline. For the 49° target the peak was on FD5 and also decreased gradually till FD15. The ANOVA indicated a significant effect of target, day, and interaction between the two (Table 1).

**Figure 2.**
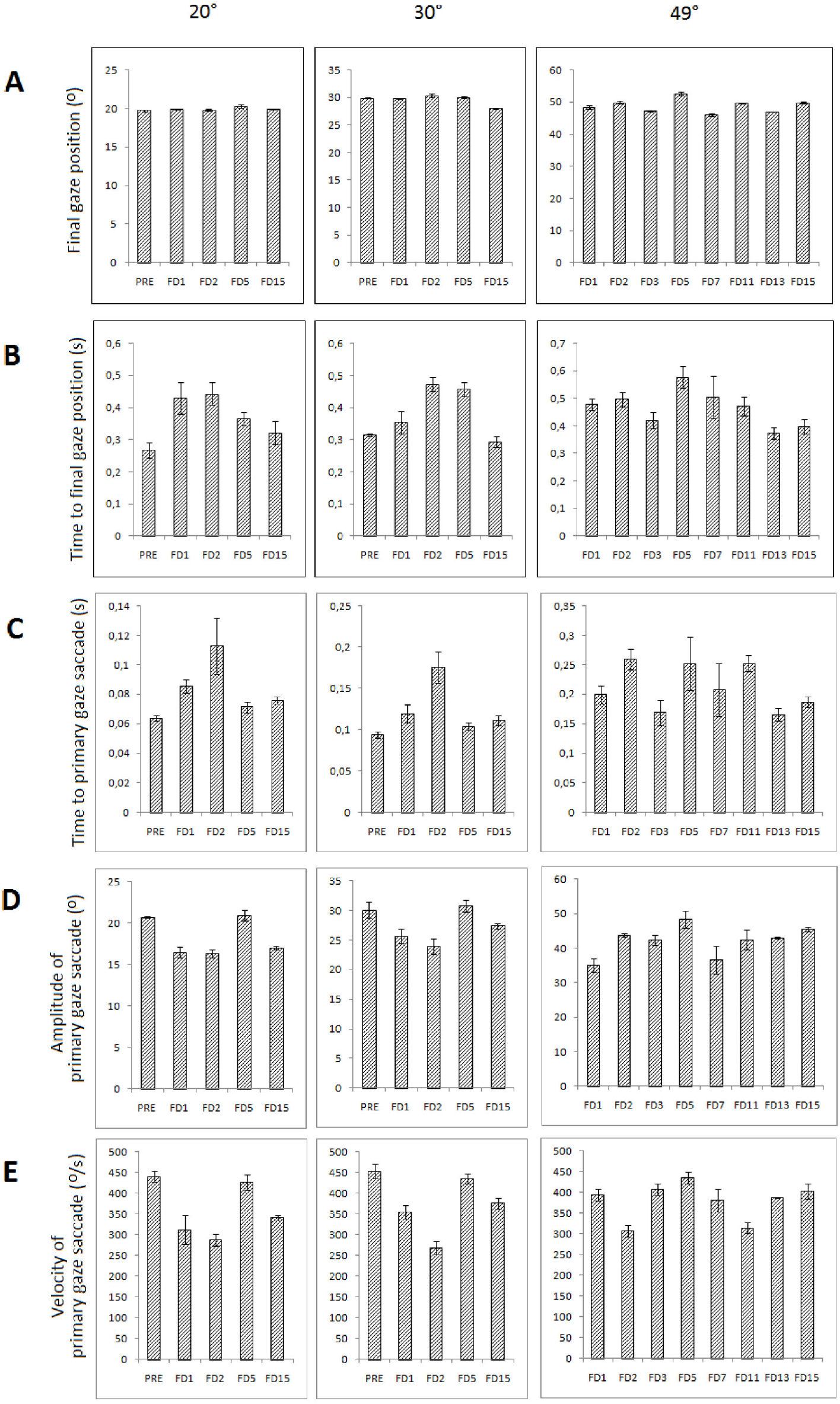
Gaze parameters. Final gaze position (A), time to primary gaze saccade (B), time to primary gaze saccade (C), amplitude of primary gaze saccade (D), and velocity of primary gaze saccade (E) as a function of flight days during target acquisition of 20°, 30°, and 49° targets. PRE: preflight, FD: flight day. Mean ± SD.

Significant changes appeared also in all other parameters as shown in Table 1 and 2, and Figure 2. The peak velocity of the primary saccade and its amplitude decreased (the peak was FD2 for the velocity to all targets; and FD2 for the amplitude to 20° and 30° targets and FD1 and FD7 to 49° target) while the time to primary gaze saccade increased (Figure 2C and D) (the peak was on FD2 to all targets).

### Head Dynamics

Since the mass of the head is larger than the mass of the eye, we expected that the change seen above in the gaze strategy during space flight were caused mainly by the change in the head strategy. As expected, for the 20° and 30° targets the time to final head position was significantly larger on FD2 compared to preflight (Figure 3A). It then decreased till FD15 but remained longer than preflight. For the 49° target the time to final head position was significantly larger on FD5 than for the subsequent flight days.

**Figure 3.**
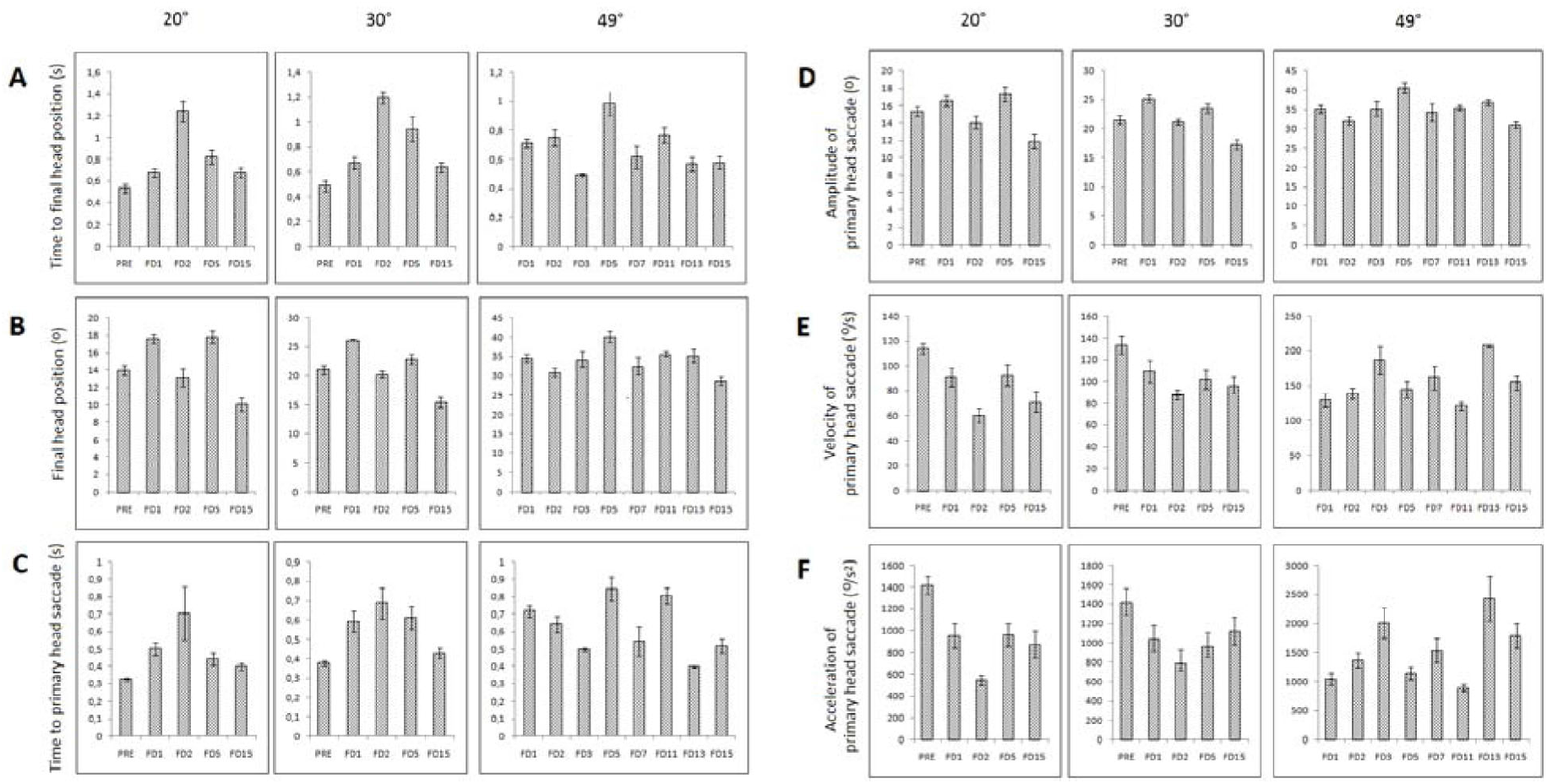
Head parameters. Time to final head position (A), final head position (B), and time to primary head saccade (C) as a function of flight days during target acquisition of 20°, 30°, and 49° targets. PRE; preflight, FD: flight day. Mean ± SD.

The final head position was significantly different across days and targets. For the 20° and 30° targets, the lowest amplitude of head movement was observed at FD15 (Figure 3B). Less changes were seen for the 49° target. For the 20° and 30° targets the time to primary head saccade was longer on FD1 and FD2 compared to preflight (Figure 3C). This duration decreased during the rest of the flight with values above the baseline. For the 49° target the maximal increase of the primary head saccade duration was on FD5 and then it decreased till FD15. The changes in the amplitude of the primary head saccade were similar to those of the final head position, i.e. significant for factors day and target, but not significant interaction between both (Figure 3D).

The changes of the peak velocity (Figure 3E) and acceleration (Figure 3F) of the primary head saccade were significant for both target eccentricity and day. The interaction was also significant for the acceleration. For the 20° and 30° targets the head peak velocity and acceleration were smaller inflight with maximal decrease on FD2. For the 49° target the peak head velocity and acceleration was smaller on FD1, FD5, and FD11 than for the other days.

### Eye Dynamics

The time to final eye position was significantly different across days and target, but there was no interaction. For the 20° and 30° targets it was the longest on FD1, retuned to baseline on FD2, and then increased again during the rest of the flight (Figure 4A). A similar decrease in time to final eye position between FD1 and FD2 was observed for the 49° target.

**Figure 4.**
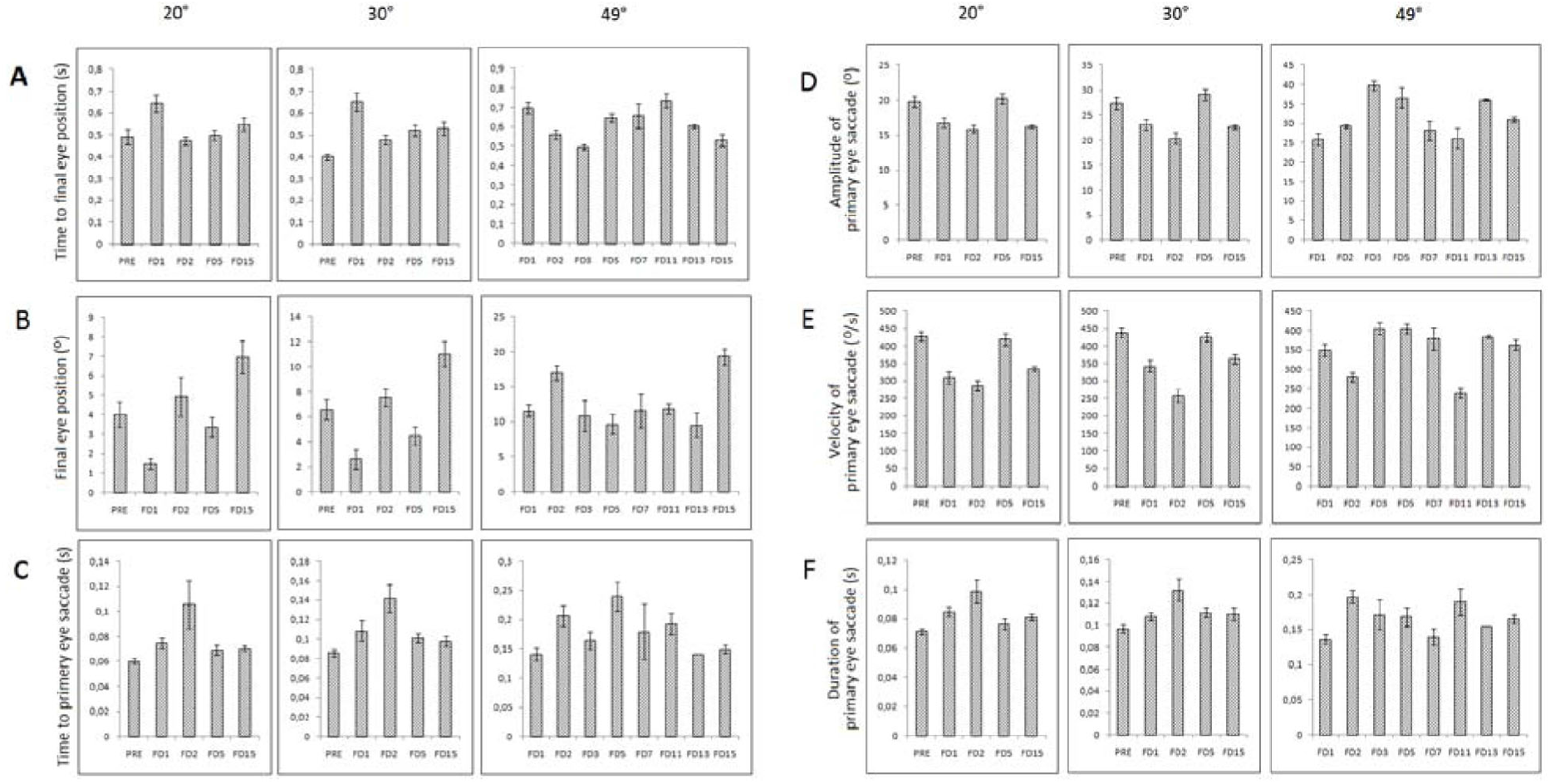
Eye parameters. Time to final eye position (A), final eye position (B), and time to primary eye saccade (C) as a function of flight days during target acquisition of 20°, 30°, and 49° targets. PRE; preflight, FD: flight day. Mean ± SD.

The final eye position was significantly different across target, day, and their interaction. For the 20° and 30° targets the final eye position decreased on FD1 compared to preflight, then returned to baseline on FD2 and decreased again on FD5 (Figure 4B). The maximal increase was on FD15.For the 49° target the final eye position was the smallest on FD5 and the largest on FD15.

The time to primary eye saccade was different across target, day, and their interaction. However, the pattern of change across flight days was not the same as for the final position time (Figure 4C). For the 20° and 30° targets the time to primary eye saccade was the longest on FD2. For the 49° target there was an increase between FD1 and FD2 similarly to 20° and 30° targets but it was the longest on FD5.

The amplitude of the primary eye saccade was significantly different across target, day, and their interaction. For the 20° and 30° targets the amplitude was smaller than preflight on FD2, returned to baseline on FD5, and then decreased again on FD15 (Figure 4D). For the 49° target, it was smaller on FD1 and FD2 compared to FD5 where was the peak increase.

The peak velocity of the primary eye saccade was significantly different across day but not across eccentricity and their interaction. For the 20° and 30° targets the peak eye velocity decreased on FD1 and FD2 compared to preflight, and returned to baseline on FD5 (Figure 4E). For the 49° target the peak eye velocity similarly decreased on FD2, compared to FD1. It was the smallest on FD2 and the highest on FD3 and FD5.

The duration of the primary eye saccade was significantly different across day, eccentricity, and their interaction. For the 20° and 30° targets the duration increased with peak on FD2 compared to baseline, and then returned to normal (Figure 4F). For the 49° target, the eye saccade duration also increased on FD2 and after that, till the end of the flight, it was relatively lower.

The correlations between head and eye movements calculated for factors day and target for the 20°, 30°, and 49° were the following:

a. Time to primary saccade: r = 0.529 (P < 0.05)
b. Amplitude of primary saccade: r = 0.639 (P < 0.05)
c. Time to final position: r = 0.294 (NS)
d. Final position: r = 0.194 (NS)
e. Peak velocity of primary saccade: r = 0.394 (NS)

## DISCUSSION

This study investigated the dynamics for gaze behavior in target acquisition in 34 astronauts participating in 20 Space Shuttle missions lasting 10-15 days. The results showed that the transition from 1 g to 0 g followed by two weeks in 0 g significantly affected eye-head coordination during the acquisition of visual targets at different angular distances. We observed an increase in the duration required to reach the target (final position), which indicates a decrease in performance. We also observed significant changes in amplitude/velocity/acceleration of the primary eye and head saccades as a function of flight day. These findings show a clear adaptation in astronauts’ gaze strategy after insertion into weightlessness.

### Saccades and Smooth Pursuit during Previous Studies

Previous studies on visual target acquisition in humans during space flight are rare and show conflicting findings. Uri et al. (1989) studied saccadic eye movements toward 10° and 20° targets in 6 astronauts during 6-8 days Space Shuttle flights. Peak eye velocity decreased in 5 subjects, but increased in 1 subject. The amplitude of the primary eye saccade tended to decrease, but this change was not found to be significant. However, only one subject was tested from FD1 to FD5; the other 4 subjects were tested only 1 or 2 days during the flight. Tomilovskaya et al. (2011) studied visual target acquisition in 7 cosmonauts before and after 6-month space flights. After landing, 3 subjects showed a decrease in peak eye velocity but no significant decrease in peak head velocity, whereas 4 subjects showed an increase in peak head velocity. The authors tried to explain the difference by the fact that some subjects were pilots and the others not. A recent report (Reschke et al. 2017) showed that the mean time to acquire visual targets after the flight was slower than before the flight, and that performance returned to baseline after 48 hours. Results were similar after space flight with duration ranging from 6 to 17 days.

Andre-Deshays et al. (1993) showed that the amplitude of the primary eye saccades to horizontal targets overshot when tested on FD5 in 1 cosmonaut, and returned to baseline on FD18 and FD22 (the primary saccades of the second cosmonaut tested only on FD20 were on target). The two cosmonauts did not show any change in vertical and horizontal smooth pursuit in orbit. By contrast, Kornilova et al. (1993) reported changes in pursuit tracking of targets moving vertically during space flight in 5 cosmonauts. Early in microgravity the pursuit was lacking behind the stimulus and corrective saccades appeared. As the flight progressed smooth pursuit deteriorated. Immediately postflight smooth pursuit improved, which the authors consider as indicating that the deficiency in microgravity may be of central origin. The authors however did not specify in which flight days their data were collected.

A decrease in corrective saccades velocity and amplitude during horizontal sinusoidal smooth pursuit was observed in 4 astronauts on flight days 2, 5, 10, and 16 (Somers et al. 2002). An immediate recovery occurred after landing, because no differences in saccade velocity and amplitude were seen postflight compared to preflight. Vesterhauge et al. (1984) found that the velocity of the saccades did not change significantly in 9 subjects during the microgravity phase of parabolic flight compared to normal gravity. Finally, experiments on monkeys in space showed gaze hypermetrics, a decrease in head amplitude and velocity, and an increase in saccadic velocity (Kozlovskaya et al. 1994, Cohen et al. 2005, Dai et al. 1998).

### Eye-Head Coordination Strategies for Gaze Performance

Our experiment performed on a larger number of days, beginning with FD1, and on a larger number of subjects than previous studies shows more comprehensive and consistent results than the previous studies above. For the 20° and 30° targets, on FD1, the time to reach the final gaze position increased compared to preflight because of a decrease in gaze velocity. This decrease in gaze velocity is caused by a decreased velocity of both eye and head primary saccades. In addition, the head movement is slower than the eye movement and, interestingly, with increased amplitude. At the beginning of the flight, the astronauts presumably reduce their head motion to mitigate space motion sickness. This slower head acceleration/deceleration (which is the stimulus that provokes motion sickness symptoms) leads to an increased duration for the head movement. On FD2 the time to reach the final gaze position also increased, presumably because of a different gaze strategy. We believe this increase is due to a further decrease of the eye and head velocity because the amplitude of the head movement decreased. The time to reach the final gaze position on FD5 and FD15 showed gradual return to baseline, which suggests an adaptation to microgravity.

Both the peak velocity and amplitude of the primary eye saccade returned to baseline earlier (on FD5) compared to head velocity/acceleration. This indicates an earlier adaptation of the saccadic movement compared to the head velocity/acceleration, which remained consistently slower than normal, presumably to minimize space motion sickness. For the 49° target the time to reach the final gaze position had its peak of increase later, on FD5, compared to short distance targets (20° and 30°). This also indicates a change in gaze strategy. After FD5, the time to reach the targets gradually reduced with a maximum on FD13 and FD15, which we assume is close to normal values, basing on the dynamics of the results of the 20° and 30° targets. We explain the target distance effect by two reasons. One is the strategy of rotation of the head with its big mass toward an approximately double angular distance. The second reason is the astronauts’ reduction of their head motion toward a larger angular distance in order to attenuate space motion sickness symptoms. These reasons are supported by the decrease in head in velocity and amplitude throughout the flight.

### Underlying Adaptive Mechanisms

The observed changes in eye-head coordination during target acquisition are presumably due to a combination of several factors and mechanisms. One is the velocity storage mechanism, which is generated in the vestibular nuclei and controlled through the nodulus and uvula of the vestibulo-cerebellum. It has been shown that in microgravity the velocity storage orientation moves from an allocentric, gravity-referenced frame to an egocentric, head-referenced frame (Dai et al. 1994, Moore et al. 2005). Another hypothesis is that humans can have several different sets of reflexes, between which they are able to switch rapidly based on the environment in which they are immersed. Procedures that help to transfer sensorimotor skills from one environment (e.g. gravitational) to another must be learned (Shelhamer & Clendaniel 2002). Factor that supposedly influences eye-head coordination in the gaze performance is a change in the perception of space. Weightlessness modifies the vestibular and somatosensory inputs, which contribute to a mental representation of 3D space in the vestibular cortex (Clément et al. 2010). A change in the mental representation of space may have an effect on the estimation of target eccentricity and distance, and therefore target acquisition performance.

Alterations in pursuit eye movements related to space flight have been attributed to changes in the tonic levels of otolith activity in the vestibulo-cerebellum (Andre-Deshays et al. 1993). Such change in tonic activity inflight could affect also gaze target acquisition. In a gravitational field, the otoliths contribute to the brain function in defining the position of the head in space by detecting head tilt with respect to the gravity vector. In the absence of a gravitational reference during space flight, the static otolith signals are ineffective and astronauts relay mainly on visual and proprioceptive information to interpret the position of the head. Therefore the altered vestibular and somatosensory inputs may lead to changes in mental representation of space (Bigelow & Agrawal, 2015). The wrong interpretation of the head’s position in space would lead to an altered perception of gaze position, which could alter the brain commands for head and eye movement during gaze target acquisition.

Other contributing factors for altered eye-head coordination in gaze performance during space flight are body fluid shifts and microcirculatory changes in brain, head and neck tissues, muscles and ligaments (Stevens et al. 2005, Schneider et al. 2008). These changes affect not only the mass of head and neck, but also the cues from the proprioceptors and motor performance including mechanically (Thornton et al. 1987). The absence of gravity causes increase in the interdiscal space in the cervical region due to absence of weight pressure from the body mass, which is another biomechanical factor that could influence target acquisition when head movement participates (Kolev & Reschke 2016).

It has been reported (Stevens et al 2005, Schneider et al 2008) that in 0 g the long-term redistribution of blood results in an increase of blood volume and blood pressure within the head. The associated increased supply of oxygen to the brain could lead to physiological changes in the central nervous system (CNS), which can affect performance (De Santo et al. 2005). In addition, it has been found that the electro-cortical activity in the frontal brain regions (Brodman areas 6 and 9) increased in weightlessness during parabolic flight (Brümmer et al. 2011). These frontal brain regions play a critical role in the planning and coordination of movements and in spatial memory. An increase in cortical activity may reflect mechanisms by which the CNS detects and processes changing gravity conditions in order to maintain motor performance. Other evidence of changes in brain cortical activity in weightlessness comes from studies showing increased beta oscillation and decreased cognitive performance (Schneider et al. 2007, Schneider et al. 2008).

Motion sickness symptoms, like somnolence, malaise and lethargy, also influence sensorimotor performance by causing delay, and producing slower, less accurate saccades (Thornton et al. 1987). Among the probable factors underlying the observed changes in eye-head coordination in the present study could be neural mechanism of functional rearrangement in the brain centers regulating patterns of motor behavior. A number of investigators have assessed the role of vestibular-based subsystems during space flight (see review in Clément & Reschke 2010). Another hypothesis is a mechanism of separation between the neural networks that control the eye and head during a coordinated action. Vasudevan & Bastian (2010) have demonstrated that there is partial overlap in the functional networks coordinating different motion. Similar findings are established in other vertebrates (McLean et al. 2008).

## Conclusion

The findings of this study are important for space exploration because the control of eye-head coordination in gaze performance is critical for spatial orientation, piloting, and locomotion in altered gravity. The current plan of space agencies is to send humans in space on long-duration missions to explore the surface of Moon and Mars. These missions will include transitions between different gravity levels (1 g, 0 g, 0.16 g, 0.38 g). It is critical to know how the brain strategies of eye-head coordination in gaze behavior adapt to these transitions for minimizing the risks of impaired control of spacecraft and the spatial disorientation due to vestibular and other sensorimotor alterations.

## ACKNOWLEDGEMENTS

This study was funded by the Extended Duration Orbiter Medical Project of the NASA Johnson Space Center Space Life Sciences Directorate (Reschke) and Centre National d’Etudes Spatiales (Clement). We would like to thank the participating crewmembers and the personnel of the Neuroscience Laboratory at the NASA Johnson Space Center for their help with data collection and data analysis.

